# Speaker responses to audience-induced social-evaluative threat: Evidence from scientific presentation tasks in immersive virtual reality

**DOI:** 10.1101/2025.09.03.673376

**Authors:** Sue Lim, Ralf Schmälzle, Gary Bente

**Affiliations:** Brian Lamb School of Communication, Purdue University, West Lafayette, 47906, IN, USA; Department of Communication, Michigan State University, East Lansing, 48824, MI, USA

**Keywords:** Public Speaking, Virtual Reality, Communication, Communication Apprehension, Social Cognition

## Abstract

Success in public speaking hinges on engaging an audience - a high-stakes social interaction that remains a significant source of anxiety and stress for many. Using a virtual-reality (VR) paradigm, we tested how speakers delivering scientific talks perceive and respond to supportive vs. unsupportive audiences. We collected behavioral (eye contact, speech rate, motion expressiveness/openness), physiological (heart rate, EEG, breathing rate, pupil dilation), and self-report measures to assess audience effects. The unsupportive audience elicited greater negative affect, arousal, and anxiety, and higher perceived cognitive and social effort. Physiologically and behaviorally, speaking to the unsupportive audience slowed the speaking rate, and acoustic analyses further indicated greater emotional arousal and vocal dominance in the unsupportive condition. Finally, VR exposure reduced speaking anxiety overall. These findings highlight VR combined with physiological measurement as a powerful approach for investigating audience effects and social-communication processes, with clear implications for augmenting social intelligence and communication skills.

## Introduction

Whether delivering a scientific talk, a business pitch, or a university lecture, public speaking is a key competency that 21^st^ century knowledge workers need to master to succeed, but also one of the most highly ranked sources of anxiety and apprehension. Public speaking is a hotbed of social communication processes (Vangelisti et al., 2013), governed by a dynamic feedback loop where a speaker’s efforts to engage an audience are met with audience reactions that can powerfully shape their performance and trigger profound anxiety. Although the term public *speaking* emphasizes verbal communication, the uniquely social nature of public speaking is perhaps even more evident in nonverbal and paralinguistic channels (Lucas and Stob, 2020). Speakers, for instance, connect with their audiences through an array of non- and paraverbal behaviors, including eye gaze, body language, or prosody.

Often overlooked, however, is the fact that audiences are also communicating with the speaker. Audience feedback occurs predominantly via social-evaluative nonverbal signals – for example, unengaged audiences make no eye contact and exhibit little positive backchannel activity (e.g., head nodding). Such social feedback signals are perceived by the speakers, which can set off a cascade of negative self-attributions, feelings and symptoms of embarrassment (e.g., blushing, dry mouth), and psychological disturbances (e.g., working memory interference, speech disfluencies (Taylor et al., 2010; Kleinlogel et al., 2020). These effects of the audience on the speaker are the topic of the current study.

This study examines the effects of the audience’s behavior on the speaker, using virtual reality as an experimental petri dish to examine social communication processes. Theoretically, our work is informed by the biopsychological (BPS) model of challenge and threat (Blascovich, 2013), which connects the social-evaluative nature of public speaking to people’s biopsychological responses. Methodologically, we leverage immersive virtual reality (VR) as well as physiological, behavioral, and subjective measures to comprehensively assessing a broad spectrum of behavioral, viscero-motor, and social-cognitive variables under realistic public speaking conditions. The results augment existing efforts in the scientific community to capture the multifaceted aspects of social processes and build better interventions targeting public speaking anxiety.

In the following, we first summarize related works and point out the research gap, which consists of causal manipulation of social factors (i.e., the audience behaviors that form the stimulus triggering social-evaluative interpretations in speakers) and the measurement of the complex dynamics underlying public speaking in the context of science communication. We then explain how the current VR-based approach offers a unique solution. Finally, we present the current study in which participants gave scientific presentations in front of two types of virtual audiences (supportive vs. unsupportive).

### Challenge, Threat, and Public Speaking Processes

The behavior of an audience is a potent form of social-evaluative feedback that can act as a significant psychological stressor, comparable to other social challenges like exclusion (Schmälzle et al., 2017) or stress interviews (Kirschbaum et al., 1993). Influential frameworks, such as the Biopsychological Model of Challenge and Threat, describe how individuals respond to such stressful performance situations (Blascovich, 2013). The model posits that individuals assess their resources and demands of a situation, leading to either a challenge or a threat response. A challenge response occurs when individuals perceive they have sufficient resources to cope with demands, resulting in adaptive physiological responses, such as increased heart rate. Conversely, a threat response arises when individuals are overwhelmed and lacking in resources, triggering negative physiological reactions, such as increased blood pressure and anxiety.

Although the challenge/threat framework was originally developed in social psychology with a focus on coping and resilience under stress, it can be applied to the public speaking situation. At the core of the challenge/threat framework lies how interpretations of challenging/threatening social situations influence physiology, with a focus on cardiovascular stress responses. While we do not directly test this model, it informs and motivates our multi-modal approach and underscores the need to simultaneously capture subjective feelings, physiological stress responses, and overt behaviors to examine public speaking in realistic settings (Sapolsky, 2004; Bradley and Lang, 2000).

This multi-modal perspective reveals the limitations of traditional approaches to study public speaking. Foundational research, particularly in work about communication apprehension, public speaking anxiety, and social stress has largely relied on observation and self-reported evaluations (e.g., Mc-Croskey, 2001, O’Brien et al., 2021). For instance, MacIntyre and colleagues provided participants with scenarios describing audience characteristics (e.g., responsiveness, pleasantness) and had them answer a questionnaire about their imagined experienced (MacIntyre and Thivierge, 1995; MacIntyre et al., 1997). Hsu (2009) had participants give impromptu speeches in front of confederate audience members and complete a questionnaire after the task. However, studies that solely use self-reports do not capture the hidden neurophysiological and overt behavioral phenomena that are ongoing as the public speaking situation unfolds, such as how speakers respond to audience reactions during their live performance.

Other researchers have examined speakers’ biological responses, such as cardiovascular reactivity and changes in heartrate, as they spoke in front of various human audiences (e.g., (Baldwin and Clevenger Jr, 1980; Hilmert et al., 2002; McKinney et al., 1983). While these studies provided more holistic picture of the public speaking processes, there remains a methodological gap. Specifically, the studies occurred in the lab with confederates as audiences, leading to low ecological validity and cost and time spent on the confederates.

### Immersive Virtual Reality (VR) as a Tool to Study Public Speaking Processes

Within this context, immersive VR technology holds great potential for experimental research in public speaking. Immersive VR enables researchers to precisely manipulate experimental variables while simulating real-life environments and situations, enhancing ecological validity of lab-based studies (Loomis et al., 1999). This capability applies to social situation and cues as well. For instance, researchers can vary audience size, engagement levels, and positive and negative audience reactions and study the effects on the speaker. When integrated with physiological and behavioral measurements such as eye-tracking and heartrate, VR-based paradigms can illustrate the cause-effect mechanisms in real-time (e.g., Schmälzle et al., 2023). These features have led to recent research that leveraged immersive VR to study the effects of virtual audiences on speakers (e.g., Artemi et al., 2025, Girondini et al., 2024, Kroczek and Mühlberger, 2023, Pertaub et al., 2002).

Building upon this body of literature, our study examines how speakers respond to socio-evaluative threat by manipulating virtual audience behavior in immersive VR (See Figure 1). Specifically, participants were asked to give a research-based presentation to supportive as well as unsupportive computer-generated audiences (within-subject design) while immersed in a virtual environment that resembled a typical academic conference venue. We focus on scientific presentation because of its multifaceted nature: Effective scientific presentation requires sustained audience engagement and breaking-down complex ideas for the audience in addition to interesting content itself (Hey, 2024). Also, scientific presentations are often connected to important outcomes such as getting a job or persuading the audience toward a specific action, making feedback signals from the audience more salient to the speaker.

**Figure 1:**
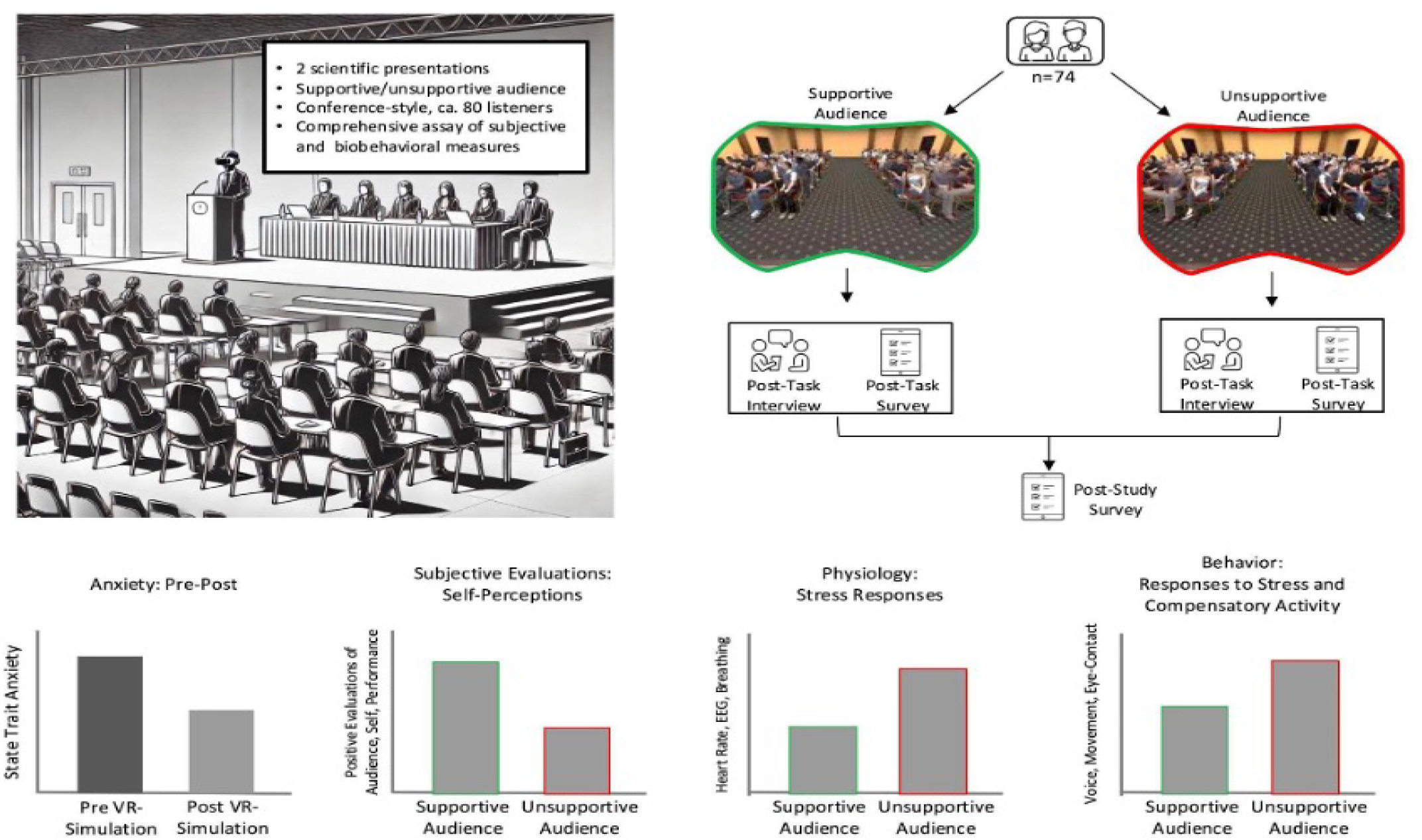
Overview of the Study Setup and Design, Main Measures, and Predictions. Participants were asked to come to the lab with two 8-12-minute scientific presentations prepared. During the study, participants wore the VR headset and gave one presentation in front of each type of audience (supportive audience vs. unsupportive audience). To comprehensively assess audience effects, we measured speaker heart rate, EEG, breathing activity, oculomotor behavior (gaze distribution and pupil dilation), body movement, and vocalics as well as subjective ratings such as experienced anxiety symptoms and physical, social, and cognitive effort required.

In terms of measurement, we combined multiple physiological (e.g., EEG, heart rate, breathing rate), behavioral (e.g., expressive motion, valence in the voice), and subjective measures (e.g., self-reported anxiety symptoms experienced during the presentation), aiming to capture not only speakers’ self-reported experience after the performance, but also their neurophysiological activity and behavior during the presentations. Our main hypotheses centered around the effect of the audience manipulation on these measures. For instance, we predicted that participants would report experiencing more anxiety-related symptoms and the pressure to invest more effort when presenting in front of the unsupportive audience. We also expected corresponding effects in other physiological measures (e.g., less eye contact with the audience, higher breathing rate). Our secondary research questions focused on the effects of the VR-based public speaking tasks on speaker anxiety and the likelihood to recommend the intervention to others.

## Results

We first present the results from the within-subject comparisons of participants’ subjective experience (see Figure 2 and Appendix Table A1). As expected, participants experienced more communication anxiety symptoms related to language and behavior (*F(1,76)* = 6.90, *p* = .010) and thought (*F(1,76)* = 16.83, *p* < .001) in front of the unsupportive audience. Furthermore, the participants felt that the unsupportive audience required more cognitive, physical, and social energy than the supportive audience (cognitive: *F(1,76)* = 11.12, *p* = .001; physical: *F(1,76)* = 5.75, *p* = .019; social: *F(1,76)* = 5.28, *p* = .024). Finally, participants reported greater levels of negative affect (*F(1,76)* = 8.09, *p* = .006) and emotional arousal (*F(1,76)* = 4.78, *p* = .032) after presenting to the unsupportive audience compared to the supportive audience. These findings demonstrate that audience’s behavior affected peoples’ social-cognitive thought patterns, even up to the level of perceptions of visceral and motor functions.

**Figure 2:**
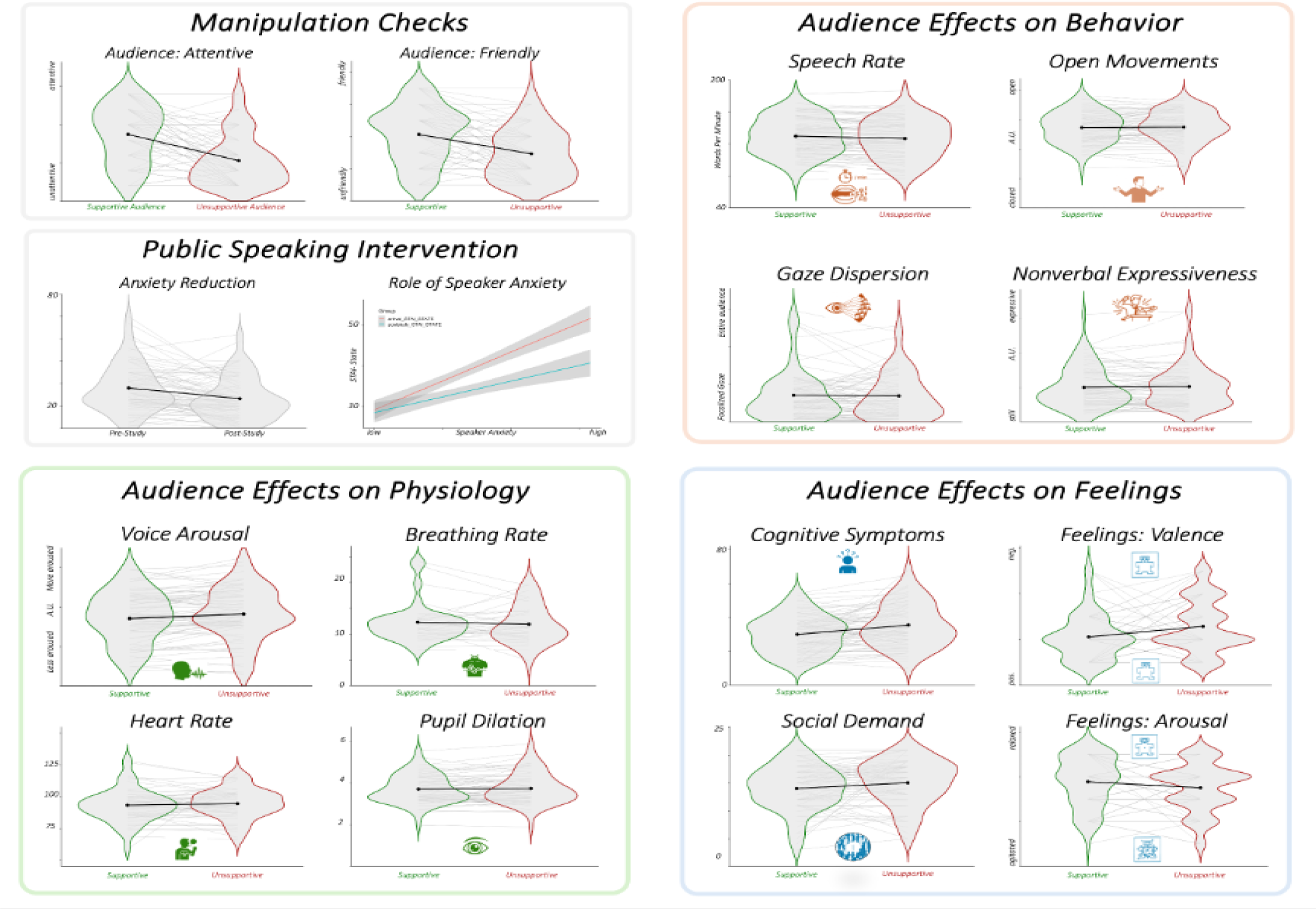
Illustration of Results: Audience Effects on Speakers’ Behavior, Physiology, and Subjective Feelings. Post-study survey data revealed that speakers were perceiving expected differences between the supportive and unsupportive audiences, in line with the intended manipulations. Results also revealed marked reductions in anxiety levels, and stronger reductions on the basis of public speaking anxiety. We assessed the effects of the manipulated audience on behavior, physiological and subjective experiences. Shown are examples of variables in each cluster. See text, tables and appendices for full results of additional variables.

**Figure 3:**
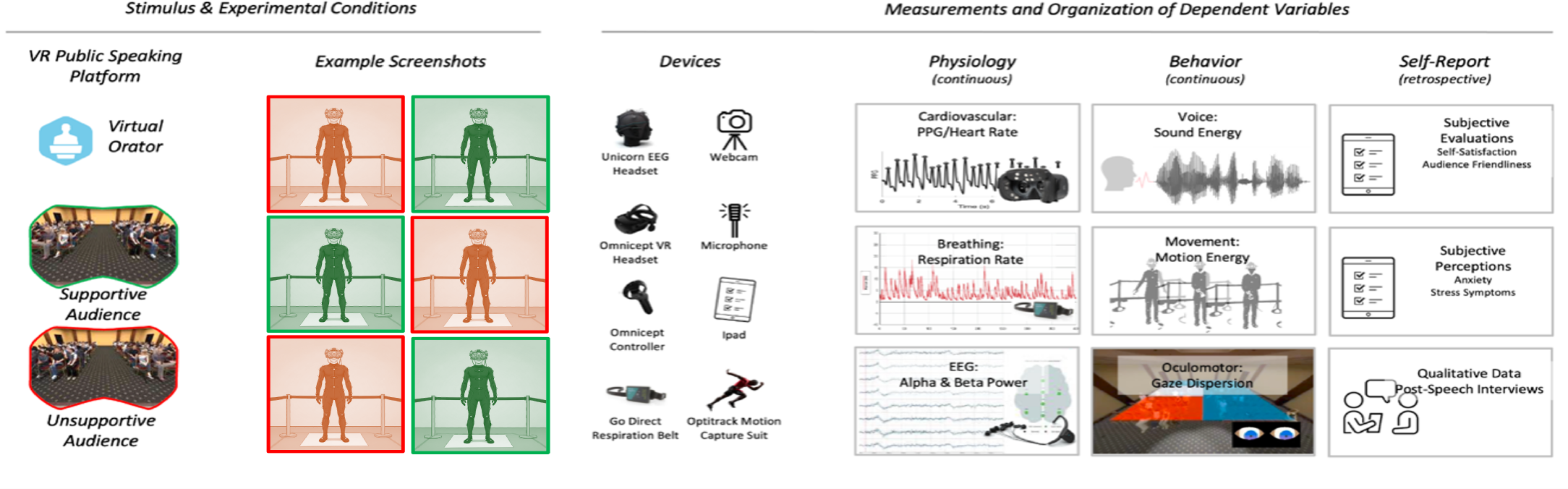
Overview of Methodology. Participants gave two scientific talks while wearing the HP Omnicept VR headset. Their physiology and behavior were recorded via the respective measurement gear, and they reported their subjective experienced through the post-presentation questionnaires. The independent variable was the type of audience (manipulated in the Virtual-Orator platform), and the dependent variables were the physiological, behavioral, and subjective responses measured, enabling a comprehensive assessment of the complex public speaking processes.

Next, we present audience effects of participants’ physiology and behavior (see Appendix Table A2). Speaking in front of the unsupportive audience decreased the words spoken per minute (*F(1,71)* = 8.11, *p* = .006) and elicited higher levels of arousal (*F(1,71)* = 12.05, *p* < .001) and dominance (*F(1,71)* = 12.05, *p* < .001) in speakers’ voices compared to the supportive audience. Additionally, speaking to the unsupportive audience also increased the alpha-beta ratio (*F(1,72)* = 4.14, *p* = .046). Other physiological and behavioral measurements did not differ by condition; however, participants who reported greater speaker anxiety prior to the intervention were less open and expressive nonverbally when presenting in front of the unsupportive vs. supportive audience (*r*_*expressiveness*_ = .28, *p*_*expressiveness*_ = .020; *r*_*openness*_ = .24, *p*_*openness*_ = .047). These results highlight how the (simulated) social audience behavior affects biological and behavioral responses, and how a speaker’s temperament or personality shapes behavioral performance under stress.

We also assessed the overall effects of the VR-based public speaking tasks as a whole. Completing the VR public speaking intervention lowered state anxiety overall (*M*_*post*_ = 33.4, *SD*_*post*_ = 9.34; *M*_*pre*_ = 38.1, *SD*_*pre*_ = 10.6; *t* = 5.67, *p* < .001), with effects more pronounced for those who reported higher levels of speaker anxiety prior to the virtual presentation (*F(1,1)* = 19.78, *p* <.001). In addition, participants were likely to recommend this VR-based public speaking simulation to others *(M* = 5.45 out of 7; *SD* = 0.54). Thus, this VR-based public speaking simulation constitutes an effective and useful communication training tool, with the added benefit that it enables experiments on how social-cognitive processes influence physiology, and behavior.

## Discussion

Here we examined how participants responded to two social-communicative challenges. While in VR, participants gave two presentations, one in front of a supportive and one in front of an unsupportive audience. We found that the virtual audience behavior strongly affected participants, demonstrating the value of VR as a social-cognitive experimentation tool, underscoring its value for training challenging public-speaking situations, and providing insights into the dynamics of speaker behavior during social-evaluative communication challenges.

First, people’s subjective experiences supported our predictions – they reported exerting greater effort and experiencing more cognitive and behavioral anxiety systems when presenting in front of the supportive audience. They also felt more negative affect and emotional arousal for the unsupportive vs. supportive audience. These results demonstrate that the experimentally manipulated virtual audience had a strong influence on participants’ subjective experience and that - despite the virtual and ultimately artificial nature of this mediated communication challenge - it produced real-world effects (e.g. Reeves and Nass, 1996).

In addition, the physiological and behavioral measures suggested that the unsupportive audience decreased the speakers’ cognitive engagement while increasing their vocal arousal and dominance. One interpretation of these results is that the unsupportive virtual audience’s behavior led speakers to compensate (e.g., try to engage the audience by sounding more excited and speaking louder), which corresponded with lower focus on the presentation content (i.e., higher alpha-to-beta ratio). While other measures did not differ significantly by condition, we found that the level of public speaking anxiety positively correlated with people’s nonverbal expressiveness and openness.

### Implications, Limitations, and Future Directions

By triangulating multiple measures – behavioral, physiological, and experiential - the current study provides a holistic picture of public speaking, one of the most important social communication skills across business, educational, and civic contexts. Methodologically, by simulating real-life communication environments and manipulating theoretical variables like audience supportiveness, we demonstrate the utility of VR-based simulations to studying complex social communication processes. This goes beyond existing work that has used VR to study people’s self-reported ratings of virtual audiences. Going forward, integrating VR-based training systems with real-time physiological and behavioral monitoring could lead to significant practical implications. For instance, future systems might monitor participants’ responses in real-time and provide adaptive feedback (e.g., virtual audience members smiling if the speaker is effectively using humor), personalized training (e.g., try “X” instead of “Y” here), or social support.

However, as with all research, this study is not without limitations. For example, in the post-experimental interview, some speakers mentioned that despite the simulation being realistic and immersive, they were aware the audiences were not real. Some speakers also mentioned that it would have been helpful if the audience behavior changed based on their performance as we suggested above future systems might. Another limitation is the occasional malfunction of some of the physiological and behavioral measurements. While the number of these malfunctions were within the range of what is expected for this type of study, it would still be desirable to have a more commodified equipment that functions reliably and with less setup and calibration effort. Lastly, most of the physiological and behavioral measures were aggregated across participants. Future work could examine individualized responding and trace dynamic changes over the duration of the presentation.

Going forward, given the rapid developments towards high-realism avatars and LLM-based agents, often combined with VR (e.g. Lim et al., 2025, Pan et al., 2025), we see a lot of potential to create more flexible and realistic audiences. For example, it would be important and feasible, to not only simulate the public speech itself, but also the Q&A session afterwards, as this session is a more bi-directional communication situation and one in which speakers are on the spot and may receive completely unanticipated questions.

### Conclusion

The current study examined how audience behavior in immersive VR impacts the speaker. demonstrating how we can use immersive VR as an experimental tool to decipher social-cognitive mechanisms, this study also highlights how VR-based experimental paradigms can be integrated with subjective, physiological, and behavioral measures to create synergy between stimulus delivery, online social cognition, and communication outcomes, and between experimental control and ecological validity. Finally, understanding social-cognitive mechanisms of public speaking holds immense practical significance. Public speaking anxiety affects many individuals and public speaking skills are a top-ranked 21st-century job skill. Therefore, leveraging VR to enhance communication skills in speakers – through training, feedback, and intervention – is a burgeoning area where insights from communication science and biology can yield tangible benefits.

## Methods

Data were collected from 80 participants (age range: 20-75; 33 Male, 42 Female, 4 Nonbinary). All participants received monetary reimbursement for their participation. Sample size was determined a-priori based on a power analysis (*α* = 0.05, 1-*β* = 0.8, paired sample *t*-test for testing the main hypotheses), which suggested that 71 participants would be required to detect a small-to-medium-sized effect (*d* = 0.3).

### Experimental Conditions and Procedures

This study employed a within-subject design in which all participants gave two presentations, one to a supportive vs unsupportive audience. The order of these presentations was counter-balanced across participants, so that half of the sample started with an unsupportive audience, the other half with the supportive audience.

For the VR-based experimental platform, we used the Virtual-Orator public speaking software (Blom, 2018). The software allows to precisely adjust audience parameters ‘audience friendliness’, ‘audience interest’, ‘audience concentration’, and ‘general distraction’. For the supportive audience, these parameters were all set to the maximum levels, resulting in audience behaviors that signaled attentiveness and friendliness to the speaker. These settings generated the attentive audience, who looked at the speaker or the projector with the slides, had an upright body posture, and neutral and friendly expressions, and did not make any distracting noise. The unsupportive audience, on the other hand, were generated by setting the respective software controls to the lowest levels, resulting in an audience that appeared distracted and unfriendly. In this condition, audience members turned away from the speaker (as if talking to the person behind them), frequently checked their cell phones, slouched, and fell asleep. Beyond audience behavior, the unsupportive condition also included loud environmental noise upon entering the room and cell phone alert and talking sounds during the presentation.

As manipulation check, we asked participants a series of audience evaluation questions in the post-presentation survey after each presentation. Specifically, participants were asked to rate the audience on 7 adjectives (e.g., attentive, friendly, cold) from 1 (not at all) to 7 (very much). The paired *t*-tests showed that participants perceived the supportive and unsupportive audiences accordingly (see Appendix Table A3), demonstrating that our manipulations worked as intended.

### Procedures

Participants filled out a pre-survey and prepared their presentation slides before coming into the lab. After providing consent to the study, we equipped the participants with the ambulatory measurements (motion capture suit, EEG, breathing belt) and the HP Omnicept VR headset and calibrated the eye-tracking system within the headset. Then, participants entered the Large-Hotel-Meetingroom-With-Chairs-environment of the Virtual-Orator public speaking platform (Blom, 2018). For each 8-12-minute presentation, we loaded the participants’ slides into the Virtual-Orator system; these slides were displayed on a virtual laptop in front of them and on the two projectors on their left and right sides. Once participants finished their presentation, they completed a post-presentation interview and survey.

### Main Measures

The measures collected in this study can be organized into three broad categories: First, over-time measures of participants’ neurophysiological and visceromotor responses, including heart rate and pupil dilation (measured via the HP Omnicept Recorder), EEG (measured via a GTec Unicorn mobile EEG system), and breathing rate (measured via a Vernier breathing belt; see Figure 2). Second, measures of participants’ behavior as they deliver the speech and respond to the audience included motion (measured via the motion-capture system), gaze behavior (generated by the Virtual-Orator platform), and their speech patterns (analyzed from presentation recordings). Third, we measured the speakers’ subjective perceptions and evaluations of the audiences as well as their experiences during the speech and global evaluations of their satisfaction with their behavior via retrospective self-report.

### Data Processing and Analysis

The processed data and the scripts are available on our Github repository (masked for review). An overview of the main derived metrics for biological and behavioral data is shown in Figure 2. In brief, all data streams were related to the common onset point - the moment participants entered the conference room. They then immediately started their speech, and the moment was marked in all separate data streams. We then extracted relevant metrics and averaged them over the first 3 minutes ^1^ of each speech (i.e. one average heart rate for the presentation in front of a supportive audience and one average HR for the presentation in front of an unsupportive audience).

Heartrate and pupil dilation metrics were directly computed by the HP Omnicept recorder software. EEG data were processed in Matlab, using the unicorn python package to read in the data, and then using EEGLab’s filtering, automated artifact correction, and power spectrum algorithms to compute global alpha and beta power spectrum values for each speech (Delorme & Makeig, 2004). Breathing rate was determined via the Vernier Graphical breathing belt analysis kit. Eye gaze was quantified via Virtual Orator’s integrated audience-zone analysis tool, which measured time spent looking at each of six audience sections (front left, front middle, front right, back left etc.). We computed the standard deviation of those metrics as a measure of equality of gaze allocation to all zones, which contained the same number of audience members. Furthermore, we used the motion-capture system to track people’s nonverbal movements throughout the speech and used python code to calculate expressiveness and openness indices. Lastly, we analyzed valence, arousal, and dominance in people’s voices using the wav2vec model (Wagner et al., 2023).

For statistical analysis of the main measures, we fitted a linear mixed effects model for each metric^2^, with audience type as the main effect variable. Intercepts varied by participant to consider the repeated-measure design, and we controlled for potential confounding effect of the speech order (i.e., which audience participants encountered first).

## Data and Code availability

Anonymized data and code is available via GitHub at https://github.com/nomcomm/NSF_Public_Speaking.

## Appendix

We present the data tables below.

**Table 1:**
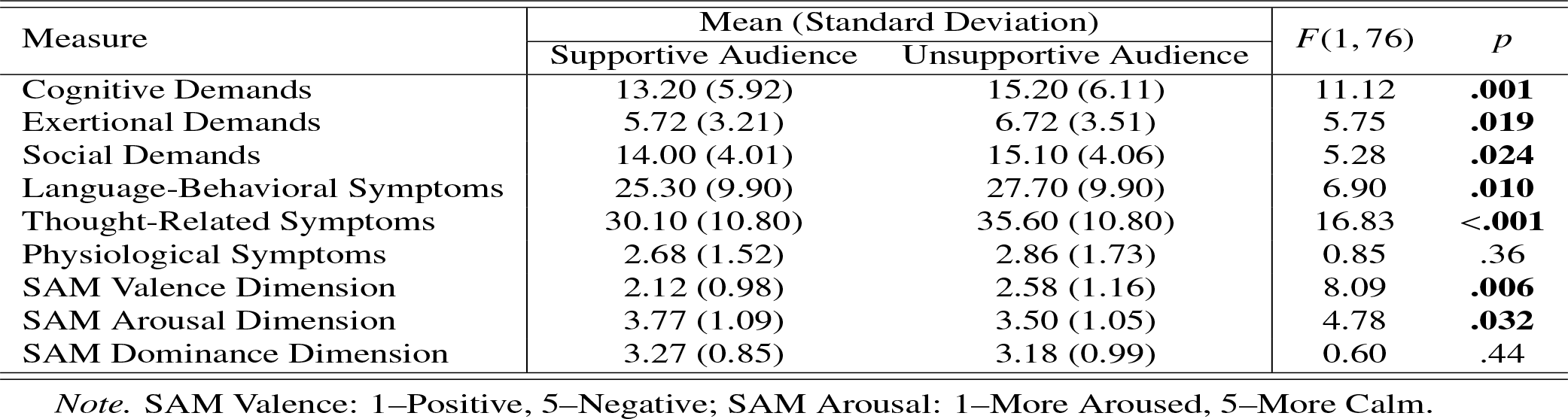
Subjective Experience by Audience Type (Supportive vs. Unsupportive)

**Table 2:** Differences in Behavior & Physiology by Audience Type (Supportive vs. Unsupportive)

| Measure | Mean (Standard Deviation) | | $df$ | $F (p\text{-value})$ |
| --- | --- | --- | --- | --- |
|  | Supportive Audience | Unsupportive Audience |  |  |
| Voice: Words per Minute | 130 (23.30) | 127 (24.60) | (1, 71) | 8.11 (**.006**) |
| EEG: Alpha-to-Beta Ratio | 6.07 (2.20) | 7.32 (4.82) | (1, 72) | 4.14 (**.046**) |
| Voice: Arousal | 0.34 (0.13) | 0.36 (0.13) | (1, 71) | 12.05 (**<.001**) |
| Voice: Dominance | 0.38 (0.12) | 0.40 (0.12) | (1, 71) | 11.30 (**.001**) |
| Voice: Valence | 0.49 (0.10) | 0.50 (0.09) | (1, 71) | 2.46 (.121) |
| Breathing per Minute (bpm) | 12.30 (3.83) | 11.90 (3.67) | (1, 31) | 0.36 (.553) |
| Motion: Openness | 68.10 (13.10) | 68.50 (13.40) | (1, 71) | 0.15 (.701) |
| Motion: Expressiveness | 2.52 (1.46) | 2.57 (1.52) | (1, 71) | 0.34 (.564) |
| Heartrate | 93.40 (13.20) | 94.70 (11.90) | (1, 52) | 1.30 (.260) |
| Pupil Dilation | 3.73 (0.68) | 3.76 (0.76) | (1, 62) | 0.31 (.579) |
| Gaze on Audience vs. Monitor* | 9.75 (24.30) | 7.82 (9.50) | (1, 67) | 0.50 (.482) |
| Gaze Dispersion* | 35.60 (33.00) | 34.70 (29.90) | (1, 74) | 0.26 (.610) |

**Table 3:** Results from Manipulation Check (Paired t-test)

| Measure | Mean (Standard Deviation) | | Effect Size<br>(Cohen's $d$) | $t (p\text{-value})$ |
| --- | --- | --- | --- | --- |
|  | Supportive Audience | Unsupportive Audience |  |  |
| Audience: Attentive | 4.37 (1.86) | 2.65 (1.65) | 0.98 | 6.90 (**<.001**) |
| Audience: Friendly | 4.13 (1.75) | 2.92 (1.57) | 0.72 | 5.48 (**<.001**) |
| Audience: Interested | 4.28 (1.76) | 2.77 (1.47) | 0.93 | 6.34 (**<.001**) |
| Audience: Lively | 3.46 (1.73) | 2.77 (1.79) | 0.39 | 3.20 (**.002**) |
| Audience: Polite | 4.74 (1.40) | 2.91 (1.50) | 1.27 | 8.25 (**<.001**) |
| Audience: Rude | 2.08 (1.28) | 3.90 (1.90) | 1.13 | -7.68 (**<.001**) |
| Audience: Cold | 3.56 (1.71) | 4.76 (1.69) | 0.70 | -4.36 (**<.001**) |

## Footnotes

1 The only exceptions to the 3-minute aggregate were gaze dispersion and screen-to-audience gaze ratio measures. These metrics were computed for the entire duration by the Virtual-Orator system. We decided on the 3-minute cut-off after examining the metrics over-time in 30 second increments. We noticed that by the 180 second mark, the metrics were generally consistent over-time, excluding certain moments of peaks and valleys. We will analyze these over-time patterns in more detail in a follow-up paper.

2 We cleaned the data before fitting the models based on the following procedure: For each metric, if the data for one speech was missing, we excluded the participant all together. This ensured adequate within-subject comparisons.

